# Rab27 GTPases regulate alpha-synuclein uptake, cell-to-cell transmission, and toxicity

**DOI:** 10.1101/2020.11.17.387449

**Authors:** Rachel Underwood, Bing Wang, Aneesh Pathak, Laura Volpicelli-Daley, Talene A. Yacoubian

## Abstract

Parkinson’s disease and Dementia with Lewy Bodies are two common neurodegenerative disorders marked by proteinaceous aggregates composed primarily of the protein α-synuclein. α-Synuclein is hypothesized to have prion-like properties, by which misfolded α-synuclein induces the pathological aggregation of endogenous α-synuclein and neuronal loss. Rab27a and Rab27b are two highly homologous Rab GTPases that regulate α-synuclein secretion, clearance, and toxicity *in vitro*. In this study, we tested the impact of Rab27a/b on the transmission of pathogenic α-synuclein. Double knockout of both Rab27 isoforms eliminated α-synuclein aggregation and neuronal toxicity in primary cultured neurons exposed to fibrillary α-synuclein. *In vivo*, Rab27 double knockout mice lacked fibril-induced α-synuclein inclusions, dopaminergic neuron loss, and behavioral deficits seen in wildtype mice with fibril-induced inclusions. Studies using AlexaFluor488-labeled α-synuclein fibrils revealed that Rab27a/b knockout prevented α-synuclein internalization without affecting bulk endocytosis. Rab27a/b knockout also blocked the cell-to-cell spread of α-synuclein pathology in multifluidic, multichambered devices. This study provides critical insight into the role of Rab GTPases in Parkinson’s disease and identifies Rab27s as key players in the progression of synucleinopathies.

## INTRODUCTION

Synucleinopathies, such as Parkinson’s Disease (PD), Dementia with Lewy Bodies (DLB) and Multiple Systems Atrophy (MSA), are a class of neurological disorders characterized by neuropathological proteinaceous inclusions consisting of α-synuclein (αsyn)(Spillantini et al., 1997). In these diseases misfolded αsyn acts as a template to induce the misfolding of endogenous αsyn(Leak et al., 2019). Targeting misfolded αsyn may lead to effective therapies to slow disease progression in these disorders, which currently have no disease-modifying therapies.

Endolysosomal pathways, including those involving Rab GTPases, have been implicated in Parkinson’s disease(Bonet-Ponce and Cookson, 2019). These pathways play roles in regulating αsyn release, endocytosis, and clearance, all of which can influence αsyn prion-like templating and inclusion formation in the brain(Bellomo et al., 2020; Malik et al., 2019; Vidyadhara et al., 2019). Rab GTPase proteins consist of a large family of >60 known mammalian members(Fukuda, 2008). Rab27 has two distinct isoforms, Rab27a and Rab27b, that work in concert to regulate a variety of processes in the endosomal-lysosomal pathway, including exosome release(Ostrowski et al., 2010), lysosomal exocytosis(Kariya et al., 2011; Shen et al., 2016), and endocytosis(Kimura et al., 2008; Kimura and Niki, 2011). In neurons, Rab27b regulates axonal transport(Arimura et al., 2009) and synaptic vesicle recycling(Pavlos et al., 2010). Therefore, Rab27 localizes to cellular pathways that play a role in αsyn regulation.

We have previously shown that Rab27b protein levels are increased in the medial temporal gyrus of PD and DLB patients(Underwood et al., 2020). Rab27b expression is elevated in the cholinergic basal forebrain of patients with Mild Cognitive Impairment or Alzheimer’s Disease and correlates with cognitive decline(Ginsberg et al., 2011). Using an *in vitro* αsyn overexpression model, we previously established that Rab27b regulates αsyn release, autophagic clearance, and toxicity(Underwood et al., 2020). To test the impact of Rab27 isoforms on fibrillary αsyn-induced aggregation and toxicity, we used the αsyn preformed fibril (PFF) mouse model in which neurons internalize fibrillar αsyn that then induces aggregation of endogenous αsyn and neuronal death(Luk et al., 2012; Osterberg et al., 2015; Volpicelli-Daley et al., 2014; Volpicelli-Daley et al., 2011).

## RESULTS

### Rab27 DKO blocks αsyn toxicity

In the PFF model, αsyn aggregation is associated with features resembling Lewy Body and Lewy neurite formation, including insolubilization and phosphorylation of endogenous αsyn at S129, and death of dopamine neurons in the substantia nigra pars compacta(Luk et al., 2012; Volpicelli-Daley et al., 2011). These effects depend on corruption of endogenous αsyn, as fibrils fail produce these phenotypes in αsyn knockout mice(Volpicelli-Daley et al., 2011). To test the impact of Rab27 in this model, primary hippocampal neurons were plated from Rab27 double knockout (DKO) or wildtype (WT) mice. Lack of Rab27a and Rab27b expression in the DKO mouse brain was confirmed by Western blot (Extended Data Fig. 1). These neurons were treated with sonicated human preformed fibrils (hPFFs) or monomeric αsyn at DIV5. αSyn aggregation was assessed by immunofluorescence against phosphorylated S129 (pS129)-αsyn, a marker for αsyn inclusion formation, at 7, 10, and 14 days after monomer or PFF treatment. αSyn monomer failed to induce pS129-αsyn staining - showing that inclusion formation depended on exposure to fibrillar αsyn (Fig. 1a). WT neurons exposed to PFFs showed abundant αsyn inclusions, while pS129-αsyn levels were dramatically decreased to background levels in Rab27 DKO neurons treated with hPFFs at all time points examined (Fig. 1a, b).

**Figure 1.**
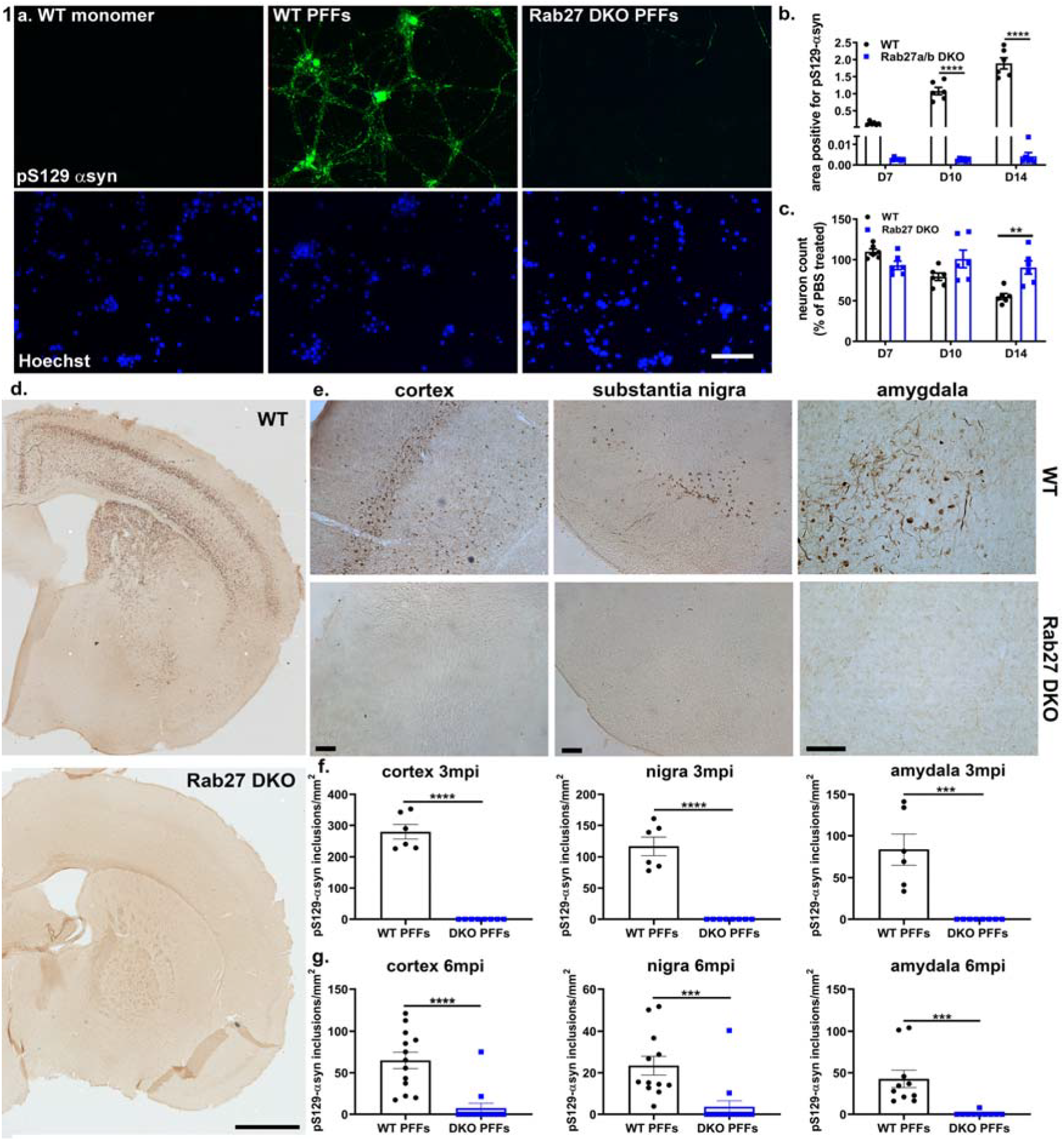
Rab27 DKO decreases αsyn aggregation. a, b) pS129-αsyn staining area in PFF- or monomer-treated WT and Rab27 DKO cultures (2-way ANOVA, Sidak’s test, n=6). Scale bar, 100 μm. c) Neuronal counts of PFF- or monomer-treated WT and Rab27 DKO cultures (2-way ANOVA, Sidak’s test, n=6). Scale bar, 1000 μm. d-g) pS129-αsyn positive inclusions at 3 mpi (f; n=6-8) and 6 mpi (g; n=10-14) in PFF-injected WT and Rab27 DKO mice in the CTX, SN, and amygdala (Student t-test). Scale bar, 100 μm. **p<0.01, ***p<0.001, ****p<0.0001. Error bars = SEM

PFF treatment induces significant neuronal death 14 days after PFF treatment, after the initiation of αsyn aggregation(Volpicelli-Daley et al., 2011). As previously observed(Volpicelli-Daley et al., 2011), a 45% loss in neuronal counts by NeuN staining was seen in WT cultures treated with hPFFs compared to those treated with monomer (Fig. 1c). In contrast, neuronal loss induced by PFFs was significantly mitigated to 10% in Rab27 DKO neurons (Fig. 1c). Together these data indicate that Rab27 DKO protects against αsyn fibril-induced aggregation and toxicity in primary neurons.

We next examined the impact of Rab27 isoforms on αsyn aggregation, neuron loss, and behavioral deficits *in vivo* using intrastriatal PFF injections to initiate αsyn aggregation and neuronal death(Luk et al., 2012; Osterberg et al., 2015). WT and Rab27 DKO mice were injected with mouse αsyn monomer or mouse PFFs (mPFFs) into the dorsolateral striatum at 8-12 weeks of age and were then examined for αsyn aggregation by pS129 αsyn immunohistochemistry (IHC) in several brain regions at 3- or 6-months post injection (mpi) (Fig. 1d-g). WT mice showed widespread inclusion formation in the cortex (CTX), substantia nigra (SN), and amygdala at 3 mpi, whereas Rab27 DKO mice showed no inclusion formation in these areas at 3 mpi (Fig. 1d-f). At 6 mpi, unlike WT mice, Rab27 DKO mice had few to no inclusions in all evaluated regions, including the CTX, SN, and amygdala (Fig. 1g; Extended Data Fig. 2). These data indicate that Rab27 DKO dramatically eliminated αsyn aggregation in the *in vivo* PFF model.

**Figure 2.**
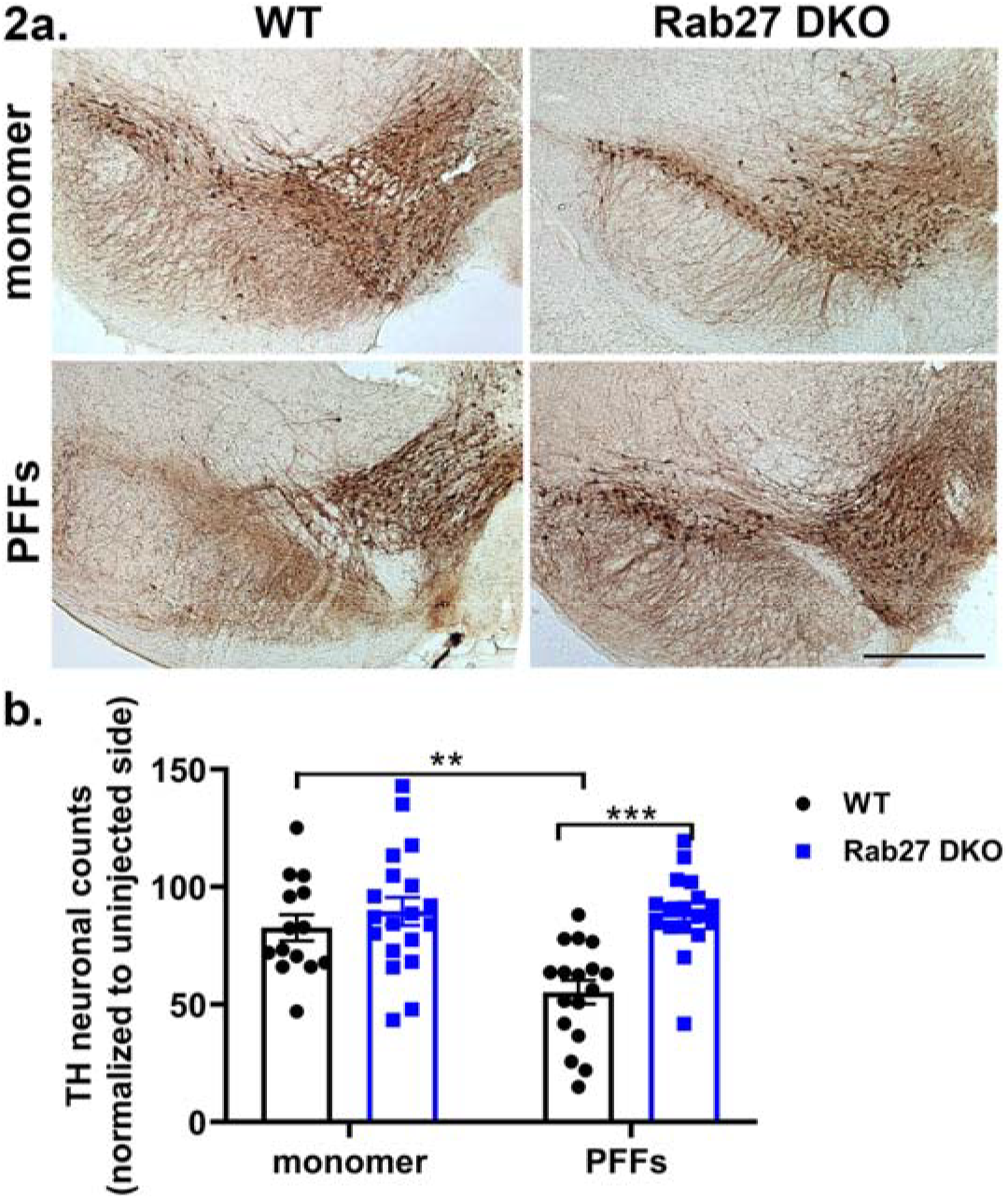
Rab27 DKO rescues PFF-induced dopaminergic neuron loss in the substantia nigra. Representative images and quantification of TH-positive cell counts analyzed by stereology in the SN pars compacta of WT and Rab27 DKO mice injected with αsyn monomer or PFFs at 6 mpi (2-way ANOVA, Tukey’s test, n=14-19) **p<0.01, ***p<0.001. Scale bar, 500 μm.

Striatal PFF injection induces loss of dopaminergic, tyrosine hydroxylase (TH)-positive neurons *in vivo*(Luk et al., 2012). To assess if Rab27 DKO rescues αsyn-induced neuronal toxicity *in vivo*, we evaluated TH-positive cell counts in the SN by stereology at 6 mpi. In WT mice, PFF injection induced a significant ~45% loss of TH-positive neurons in the ipsilateral SN compared to the contralateral SN at 6 mpi, as previously described(Luk et al., 2012) (Fig. 2). Rab27 DKO prevented TH-positive cell loss by mPFFs compared to WT mice injected with mPFFs (Fig. 2).

The elimination of αsyn aggregation and neuronal loss in Rab27 DKO mouse brains was associated with a rescue of behavioral deficits. At 3 mpi, PFF-injected Rab27 DKO mice demonstrated significantly better motor function on the pole test compared to WT mice injected with PFFs (Fig. 3a). Although studies vary in their reports of motor deficits in the PFF model likely dependent on behavioral paradigm differences(Hayakawa et al., 2020; Luk et al., 2012; Stoyka et al., 2020), we did not observe a consistent PFF-induced motor deficit on either the pole test or rotarod tests at 6 mpi (Fig. 3b, Extended Data Fig. 3d). Of note, Rab27 DKO mice demonstrated a slight but significant decrease in latency to fall on the rotarod test at 6 mpi in both PFF- and monomer-injected mice (Extended Data Fig. 3d). These genotypic effects could be due to the increased weight of Rab27 DKO mice compared to WT mice (Extended Data Fig. 3f, g).

**Figure 3.**
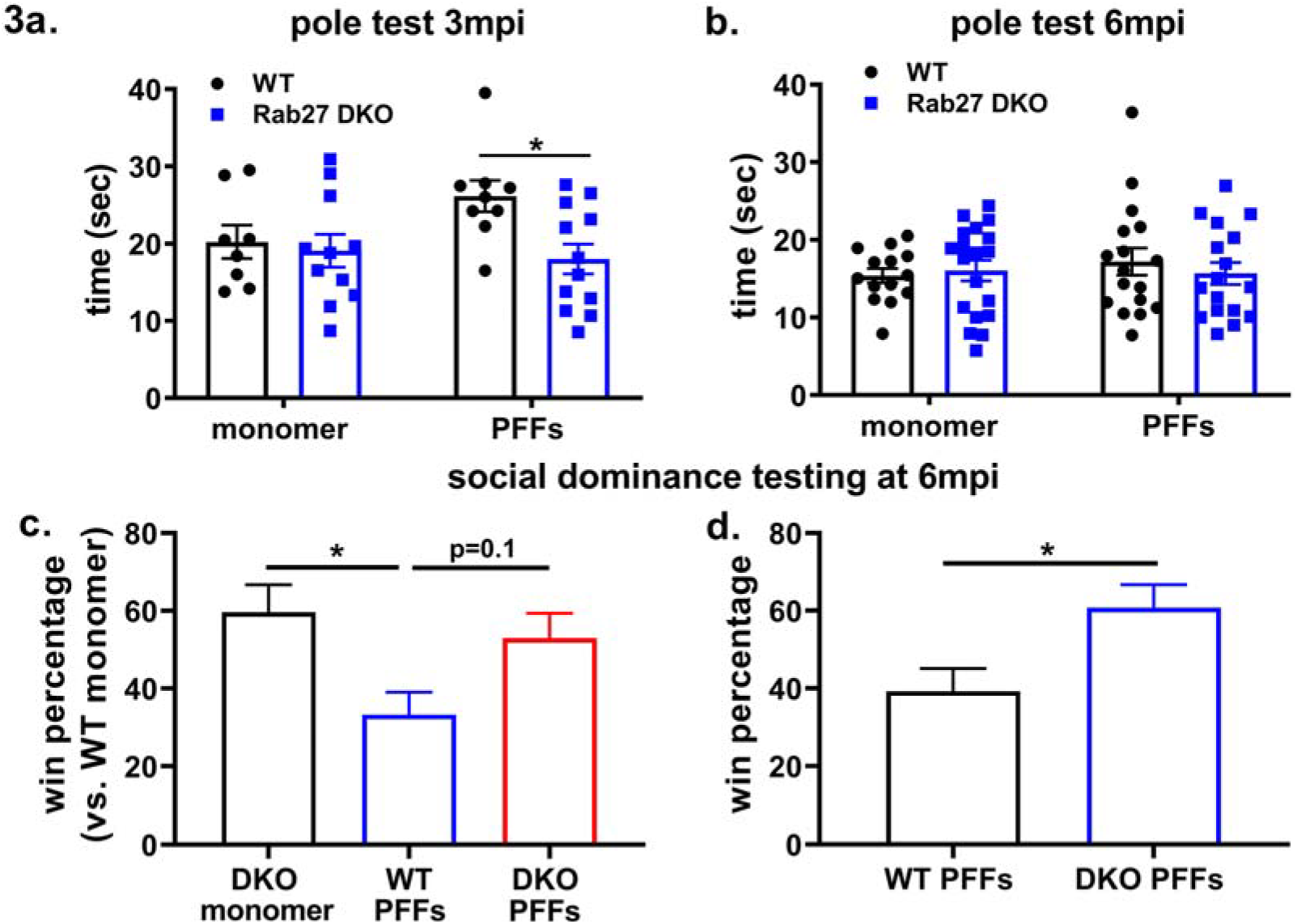
Rab27 DKO rescues behavioral deficits induced by PFF injection. a, b) Time to descend on the pole test of WT and Rab27 DKO mice injected with αsyn monomer and PFFs at 3 mpi (a; n=8-12) and 6 mpi (b; n=14-19) (2-way ANOVA, Tukey’s test). c) Win rates of PFF-injected WT and Rab27 DKO mice and monomer-injected Rab27 DKO matched against monomer-injected WT controls in the social dominance test (2-way ANOVA, Tukey’s test, n=14-19). d) Win rates of PFF-injected WT and Rab27 DKO mice matched against each other in the social dominance test (Student’s t-test, n=17). *p<0.05.

Non-motor deficits in the PFF model have been previously demonstrated(Stoyka et al., 2020). Similarly, we observed that WT mice injected with mPFFs showed a deficit in social dominance in the tube test, a test of prefrontal cortex and amygdala function(Arrant et al., 2016; Miczek et al., 1974; Soumiya et al., 2016; Zhou et al., 2018), at 6 mpi but not at 3 mpi (Fig. 3c, Extended Data Fig. 3e). This deficit was rescued by Rab27 DKO (Fig. 3c, d). PFF injection did not induce anxiety deficits, as measured on the elevated zero maze (EZM) at 6 mpi (Extended Data Fig. 3c).

### Rab27 DKO decreases αsyn internalization

We hypothesized that the near elimination of αsyn aggregation and neuronal toxicity in Rab27 DKO mice was due to a defect in internalization of exogenously applied PFFs. To test this hypothesis, we utilized two different assays to measure the internalization of αsyn fibrils. We first treated primary neurons from Rab27 DKO or WT mice with unlabeled hPFFs at DIV5 and performed two-step staining 14 days later to label extracellular and intracellular αsyn as previously described(Volpicelli-Daley et al., 2011). While we observed a dramatic 60% reduction in the FITC intracellular αsyn signal in Rab27 DKO neurons compared to WT neurons treated with PFFs, there was no difference between the TRITC αsyn signal, which reflects primarily extracellular αsyn (Fig. 4b). This finding points to a reduction in αsyn fibril internalization in the absence of Rab27.

**Figure 4.**
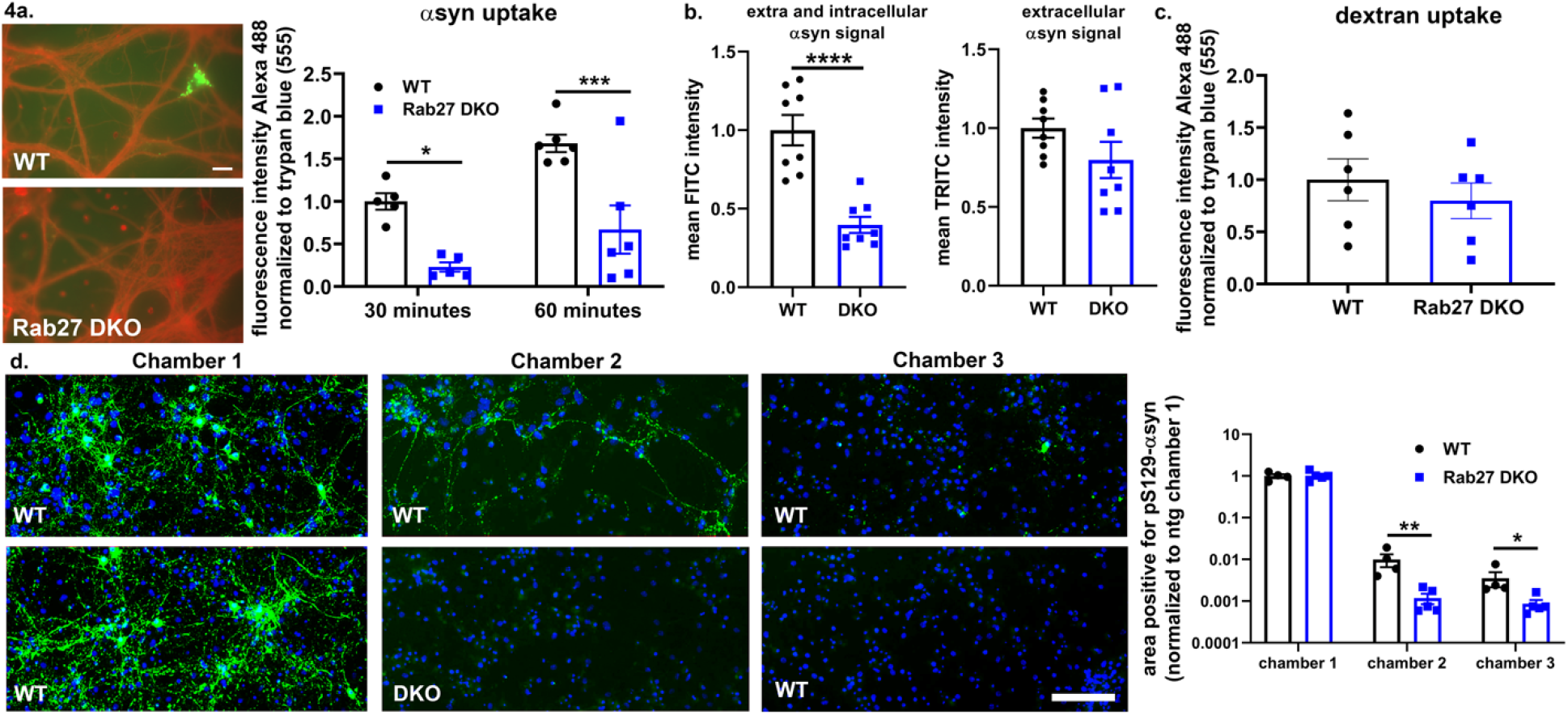
Rab27 DKO reduces αsyn internalization and cell-to-cell transmission. a) A488-tagged PFFs uptake in WT and Rab27 DKO neurons (2-way ANOVA, Tukey’s test, n=5-6). Scale bar, 20 μm. b) Intracellular and extracellular αsyn levels 14 days post PFF treatment (Student’s t-test, n=8). c) A488-dextran uptake in WT and Rab27 DKO neurons (Student’s t-test, n=6). d) pS129-αsyn in multichamber microfluidic devices. WT neurons were plated in chambers 1 and 3, and WT or Rab27 DKO in chamber 2 (Multiple t-test, Holm-Sidak correction, n=4-5). Scale bar, 100 μm. *p<0.05, **p<0.01 ***p<0.001, ****p<0.0001.

To validate these findings, we also performed live imaging of Rab27 DKO or WT neurons treated with Alexa488-labeled hPFFs. At 30 or 60 minutes following hPFF treatment, the signal of labeled hPFFs bound extracellularly was quenched by treatment with trypan-blue, and internalization of Alexa488-tagged PFFs was evaluated by live cell imaging as previously described(Volpicelli-Daley et al., 2011). Rab27 DKO neurons demonstrated a 77% decrease in A488-hPPF uptake at 30 minutes post treatment compared to WT neurons (Fig. 4a). While PFF uptake by Rab27 DKO neurons did increase somewhat at 60 minutes compared to that at 30 minutes, the amount of tagged hPFF uptake was still significantly reduced compared to WT neurons (Fig. 4a).

To test the effects of Rab27 DKO on activity-dependent bulk endocytosis, we performed live cell imaging of Rab27 DKO or WT neurons treated with A488-tagged dextran (10,000 MW) and evaluated dextran signal after washing. We found no difference in the fluorescent signal between Rab27 DKO and WT neurons, indicating no difference in the internalization of dextran (Fig. 4c).

### Rab27 DKO blocks cell-to-cell αsyn transfer

We next used microfluidic culture device system to test the impact of Rab27 DKO on cell-to-cell transfer of αsyn. These devices allow for the synaptic connections between neurons cultured in three separate compartments connected by microgrooves but prevent free diffusion of PFFs between chambers due to hydrostatic pressure differences between the compartments(Park et al., 2006). We plated WT neurons in the first and third compartments and either Rab27 DKO or WT neurons in the second compartment. After treating neurons in the first compartment with PFFs at DIV5, pS129-αsyn staining was detectable 14 days later in all three WT chambers when devices were plated with WT neurons in the second chamber, with the highest levels of staining in the first chamber and decreasing in intensity moving from the second to third chambers. Plating of Rab27 DKO neurons in the second chamber decreased pS129-αsyn staining to nearly undetectable levels in the second and third chambers, yet pS129-αsyn staining was high in WT neurons plated in the first chamber (Fig. 4d). Together these data indicate that Rab27 DKO interrupts the internalization of αsyn and its downstream neuronal propagation.

## DISCUSSION

Our data demonstrates that Rab27 isoforms regulate the uptake, cell-to-cell transmission, aggregation, and neuronal toxicity by αsyn fibrils. Rab27 DKO dramatically eliminated αsyn aggregation induced by αsyn PFFs in primary neurons and *in vivo*. This reduction in αsyn inclusion formation was associated with a reversal of behavioral deficits *in vivo* and protection against neuronal loss both *in vitro* and *in vivo*. *In vivo* Rab27 DKO resulted in the complete elimination of αsyn inclusion formation in all involved brain regions at 3 mpi. This effect persisted at 6 mpi, indicating that the elimination of αsyn aggregation was not likely due to a delay in their formation. We also demonstrated that Rab27 DKO greatly reduced the internalization of αsyn fibrils in primary neurons, pointing to a likely mechanism by which Rab27 isoforms facilitate αsyn aggregation and toxicity. While studies have pointed to other molecular targets that regulate αsyn internalization, including Myosin VIIb(Zhang et al., 2020), Lag3(Mao et al., 2016), Neurexin 1(Mao et al., 2016; Shrivastava et al., 2015), and Heparan sulfate proteoglycans(Holmes et al., 2013), the near elimination of αsyn aggregation and the prevention of neuronal loss points to Rab27 being a master regulator of αsyn fibril uptake and propagation.

Rab27s regulate endocytosis in exocytic pancreatic cell lines via its interaction with coronin 3(Kimura et al., 2008; Yamaoka et al., 2016), an actin-bundling protein that supports invagination for clathrin-dependent and clathrin-independent endocytosis(Mooren et al., 2012). Our data suggests that Rab27 isoforms regulates the endocytosis of larger molecular weight αsyn species in neurons. Rab27b has been previously shown to regulate synaptic vesicle exocytosis/endocytosis in conjunction with Rab3 isoforms at synaptic terminals(Pavlos et al., 2010). At the synapse, Rab3 isoforms promote synaptic vesicle fusion along with Rab27b, but Rab27b remains bound to the plasma membrane in its GDP-bound form and helps to facilitate synaptic vesicle recycling(Pavlos et al., 2010). We hypothesize that αsyn fibrils could become incidentally internalized during normal synaptic vesicle exocytosis-endocytosis cycles at the synapse. Extracellular, misfolded αsyn that may develop during pathological states could bind to the synaptic membrane and thus become incidentally internalized via Rab27, allowing for the propagation of pathogenic forms of αsyn.

While the reduction of αsyn uptake is likely a significant contributor to the lack of αsyn aggregation and toxicity in our studies, Rab27 DKO may also affect other cellular processes to reduce αsyn toxicity. Rab27 regulates axonal transport of TrkB and interacts with kinesin(Arimura et al., 2009). Rab27 also regulates actin transport bidirectionally by its interactions with myosin Va and myosin VIIa(Desnos et al., 2003; El-Amraoui et al., 2002; Fukuda et al., 2002; Kuroda and Fukuda, 2005; Lopes et al., 2007; Ramalho et al., 2009). αSyn oligomers bind to kinesin and block the transport of other proteins along microtubules(Prots et al., 2013). Rab27 DKO may disrupt the axonal transport of αsyn, inhibiting its ability to move throughout the neuron. The inhibition of αsyn movement may inhibit its migration from its site of endocytosis and thereby limit its ability to interact with endogenous αsyn, thus resulting in decreased aggregation potential.

The relative contribution of Rab27a and Rab27b isoforms in these effects in the fibril model is unclear. Rab27b is highly expressed in neurons in the cortex, striatum, and midbrain, areas affected in PD and DLB, while little Rab27a expression has been detected in neurons(Gomi et al., 2007; Yi et al., 2002). Given that Rab27b is the main isoform expressed in neurons, it is likely that Rab27b is the critical isoform implicated in these findings; however, as Rab27a is expressed in glial cells in the brain(Gomi et al., 2007; Yi et al., 2002), Rab27a could contribute indirectly to these effects. Since Rab27a and Rab27b share most, if not all, effector proteins(Fukuda, 2013), we used Rab27 DKO mice in order to assess the effects of Rab27a and Rab27b without the potential rescue of one homolog over the other.

We previously established that Rab27b expression is elevated in the post-mortem brain lysates of both PD and DLB patients(Underwood et al., 2020). Based on our previous data showing that Rab27b regulates αsyn clearance combined with our data presented here, we propose the following model for the role of Rab27 isoforms in synucleinopathies. Early in disease, upregulation of Rab27 serves as a compensatory mechanism in neurons to clear misfolded αsyn and prevent its intracellular buildup and toxicity. Any low levels of any endocytosed extracellular αsyn can be directed to degradation pathways. However, as disease progresses and extracellular, pathogenic αsyn levels increase, Rab27’s normal role in synaptic vesicle endocytosis becomes maladaptive, overcoming any protective function of Rab27 in protein degradation, and thus aids αsyn cell-to-cell transmission. αSyn can become internalized by neurons during synaptic vesicle recycling that is promoted by Rab27 at the synapse. As different Rab27 effectors are likely to mediate these different Rab27 trafficking functions, future directions will explore the effectors by which Rab27 impacts these various cellular functions. By focusing on the downstream targets of Rab27, it is feasible to upregulate processes that encourage the autophagic clearance of αsyn but do not promote its endocytosis, thereby reducing the negative effects of Rab27 on αsyn propagation. In conclusion, we demonstrate that Rab27 isoforms are critical regulators of αsyn uptake, aggregation, and toxicity in fibrillar models of αsyn.

## METHODS

### CONTACT FOR REAGENT AND RESOURCE SHARING

Further information and requests for resources and reagents should be directed to and will be fulfilled by the Lead Contact, Talene Yacoubian.

### EXPERIMENTAL MODEL AND SUBJECT DETAILS

#### Mice

Mice were used in accordance with the guidelines of the National Institute of Health (NIH) and University of Alabama at Birmingham (UAB) Institutional Animal Care and Use Committee (IACUC). Animal work performed in this study was approved by UAB’s IACUC. Transgenic mice with Rab27 DKO (Rab27a ^ash/ash^ Rab27b ^−/−^) were generated by Dr. Miguel Seabra at Imperial College and obtained from Dr. Brian Rudd at Cornell University(Tolmachova et al., 2007). Rab27 DKO homozygous mice were bred with C57BL/6J mice from Jackson Laboratories (Bar Harbor, ME; Cat#000664, 140 RRID:IMSR_JAX:000664) to generate Rab27 DKO heterozygous mice (Rab27a ^+/ash^ Rab27b ^+/−^). Rab27 DKO heterozygous mice were crossed together every 2-3 generations to generate back-crossed WT mice to control for genetic drift between Rab27 DKO and WT during homozygous breeding.

#### Primary Neuronal Culture

Primary hippocampal cultures were prepared from male and female postnatal 0 (P0) or P1 mice as previously described(Wang et al., 2018). For PFF experiments, neurons underwent a complete media change and were treated with PBS, monomeric human αsyn, or human αsyn PFFs (1 μg/ml) in Neurobasal-A media with B-27 supplement at DIV5. For uptake assays, primary neurons were treated with A488-tagged αsyn PFFs (1 μg/ml). A488 signal was imaged by live cell imaging at 30 and 60 minutes post treatment. Similarly, for determining the uptake of dextran, primary neurons were treated with A488-tagged dextran (100 μg/ml). A488 signal was imaged by live cell imaging at 30 minutes post PFF treatment. To confirm the live uptake assay findings, primary neurons were treated with untagged human αsyn PFFs at DIV5, and internalization of human αsyn PFFs was evaluated by two-step staining 14 days later to label extracellular and intracellular αsyn as previously described(Volpicelli-Daley et al., 2011).

#### Microfluidic chambers

Primary neurons were plated in microfluidic chambers with three cell compartments (#TCND1000, Xona Microfluidics, Temecula, CA) per manufacturer’s instructions. At DIV5, 2 μg/ml PFFs were added to chamber 1 only. A 75 μl difference in media volume was maintained between each chamber with the lowest volume in chamber 1 to control the direction of media flow.

#### Generation of α-synuclein fibrils

Recombinant human and mouse αsyn was generated as previously described(Giasson et al., 2001; Volpicelli-Daley et al., 2014; Volpicelli-Daley et al., 2011). For primary cultures, fibrils were generated by incubating purified human monomeric αsyn at a concentration of 5mg/ml. Sonicated αsyn fibrils were added to primary neuronal cultures at 1 μg/ml in neuronal media. For stereotaxic injection in mice, fibrils were generated by incubating purified mouse monomeric αsyn at a concentration of 5mg/ml. Sonicated αsyn fibrils were analyzed by dynamic light scattering on a DynaPro NanoStar (Wyatt Technology, Santa Barbara CA) to ensure sonicated fibril average radius was between 10-100 nm prior to injection.

#### Stereotaxic Injections

Rab27 DKO or C57BL/6 mice at 8-12 weeks of age were deeply anesthetized with 5% isoflurane and maintained at 0.25%-4% during surgery. Mice were unilaterally injected with 5 μg of αsyn fibrils or monomer (2 μl of 2.5 mg/ml) at a flow rate of 0.250 μl/minute into the dorsolateral striatum [anteroposterior (AP): 0.2 mm from bregma; mediolateral (ML): −2 mm from midline; dorsoventral (DV): −2.6 mm below dura] according to a previously established protocol(Luk et al., 2012). Mice were given buprenorphine (1mg/kg) before surgery and the day after to minimize pain and discomfort.

#### Immunohistochemistry and Immunofluorescence

Mice were perfused with PBS followed by 4% paraformaldehyde. After dissection, brains were sliced by microtome in 40 μm thick coronal sections. Every 6^th^ section from the anterior brain (including the sensorimotor and striatal regions) was stained for pS129 αsyn, while every 4^th^ section from the posterior sections (including the substantia nigra) was stained for pS129 αsyn and TH. Sections were imaged using an Olympus BX51 epifluorescence microscope. For pS129-αsyn positive inclusion counts, three sections in either the sensorimotor cortical regions (Bregma +1.54 mm to +0.62 mm) or the substantia nigra (Bregma −2.70 mm to −3.88 mm) per well were randomly selected and quantitated using ImageJ with the rater blind to experimental conditions. One section was selected in the central and basolateral amygdala region (Bregma −0.82 mm to −1.34 mm) and quantitated using ImageJ with the rater blind to experimental conditions. αSyn inclusion counts were normalized per mm^2^ area.

#### Stereology

Stereological estimates of TH-positive neuronal numbers were performed using the optical fractionator method of the StereoInvestigator 8.0 software from MBF Biosciences (Microbrightfield Inc., Williston, VT, USA) as previously described(Ding et al., 2015). Stereology estimates were done with the investigator blinded to the experimental condition.

#### Behavior

Behavior tests to assess motor, anxiety, and social functions were conducted 3 and 6 months after PFF injection. Mice were handled for 3 to 5 days before testing began and habituated to testing room for 30 minutes at the start of each testing day. The researcher conducting the tests was blinded to injection type but not to genotype due to the Rab27 KO *ashen* silver coat phenotype. Mice were evaluated by elevated zero maze, open field, social dominance tube test, pole test, and rotarod. Details of these behavioral tests are in the Supplemental Materials.

#### Statistical Analysis

GraphPad Prism 8 (La Jolla, CA) was used for statistical analysis of experiments. Data were analyzed by either Student’s t-test, one-way ANOVA, or two-way ANOVA, followed by post-hoc pairwise comparisons using Sidak’s or Tukey’s multiple comparison tests. Statistical significance was set at p ≤ 0.05. All data values are presented as mean ± SEM. Statistical analysis details, including F-statistics, t-statistics, p values, and sample sizes, for each experiment are found in the Supplemental Materials.

#### Reporting Summary

Further information regarding the experimental design is found in the Research Reporting Summary linked to this paper.

#### Data Availability

Source data are available for graphs plotted in Figures 1–4 and Extended Data Figures 1–3. Scans of full western blots can be found in Supplemental Figure. All other data are available from the corresponding author upon reasonable request.

## ACKNOWLEDGEMENTS

This study was supported by NIH [R01 NS088533 (TAY); R01 NS112203 (TAY); R56 NS115767 (TAY); P50 NS108675 (TAY); F31 NS106733 (RU)], American Parkinson Disease Association, and the Parkinson Association of Alabama. We thank Dr. Miguel Seabra at Imperial College and Dr. Brian Rudd at Cornell University who provided the Rab27 DKO mouse line that was used in these studies.

## AUTHOR CONTRIBUTIONS

RU designed and performed experiments, analyzed data, and wrote manuscript. BW designed and performed experiments, analyzed data, and reviewed manuscript. AP performed experiments and reviewed manuscript. LVD provided reagents, assisted in uptake experiments, and edited manuscript. TY designed experiments, analyzed data, and wrote and finalized manuscript. All authors read and approved the final manuscript.

## COMPETING INTEREST DECLARATION

The authors have no competing interests to declare.

## ADDITIONAL INFORMATION

Supplemental Information regarding methods, detailed statistical analysis, and full Western blot data is available for this paper. Correspondence and request for materials should be addressed to Talene Yacoubian.

## EXTENDED DATA FIGURES

**Extended Data Figure 1.**
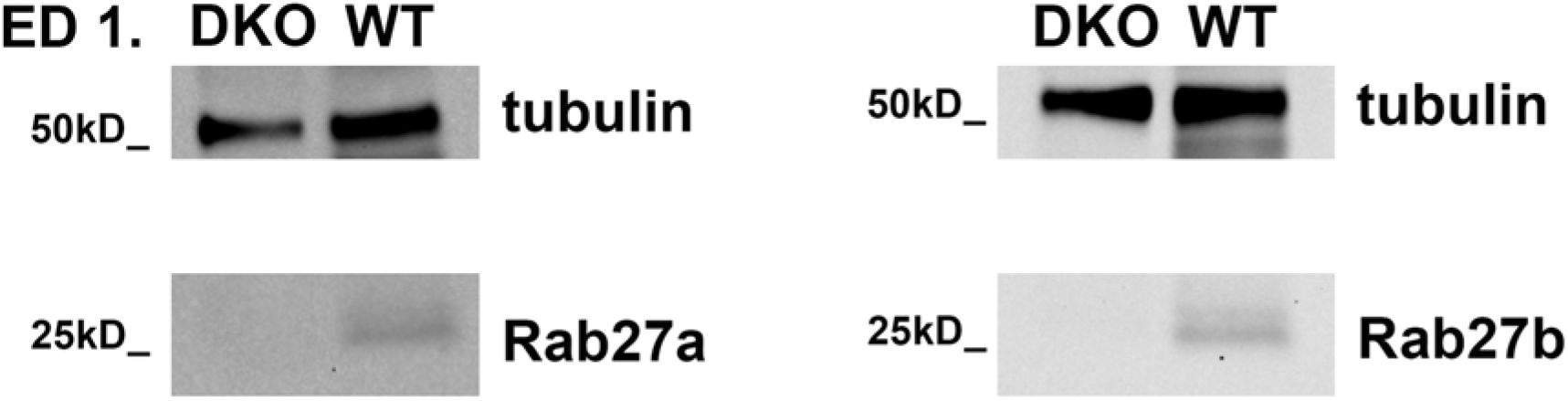
Rab27a and Rab27b are not expressed in the brain lysates of Rab27 DKO mice. Rab27 DKO was confirmed by Western blot for Rab27a and Rab27b expression in WT and Rab27 DKO mouse cortical brain lysates.

**Extended Data Figure 2.**
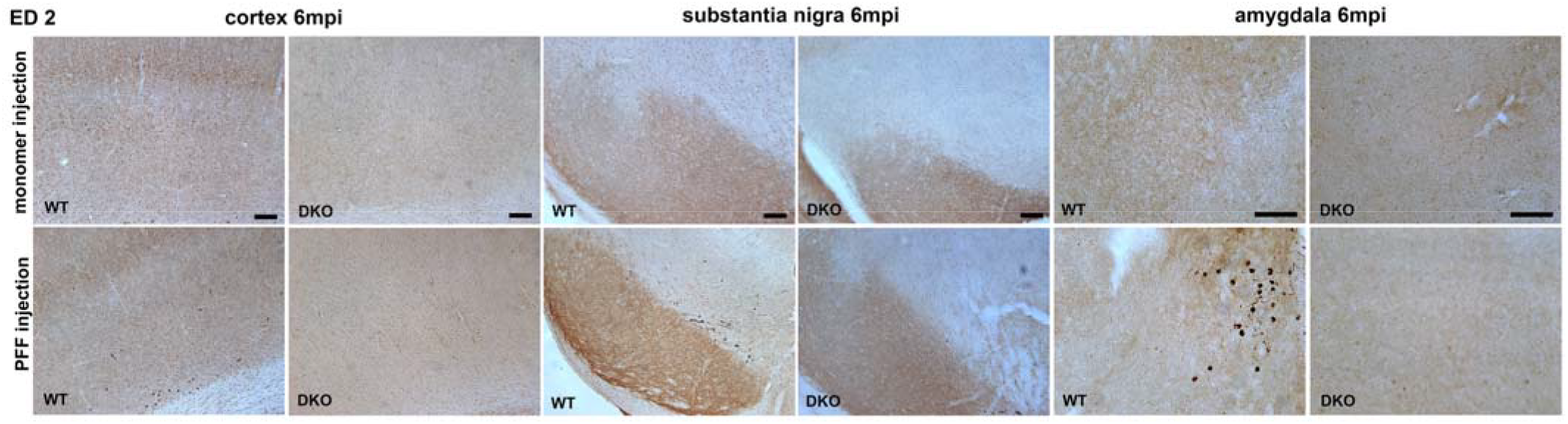
Rab27 DKO reduces αsyn aggregates at 6 months after PFF injection. Representative images of pS129-αsyn-positive inclusions in monomer- or PFF-injected WT and Rab27 DKO mice at 6 mpi in the CTX, SN, and amygdala. Scale bar, 100 μm.

**Extended Data Figure 3.**
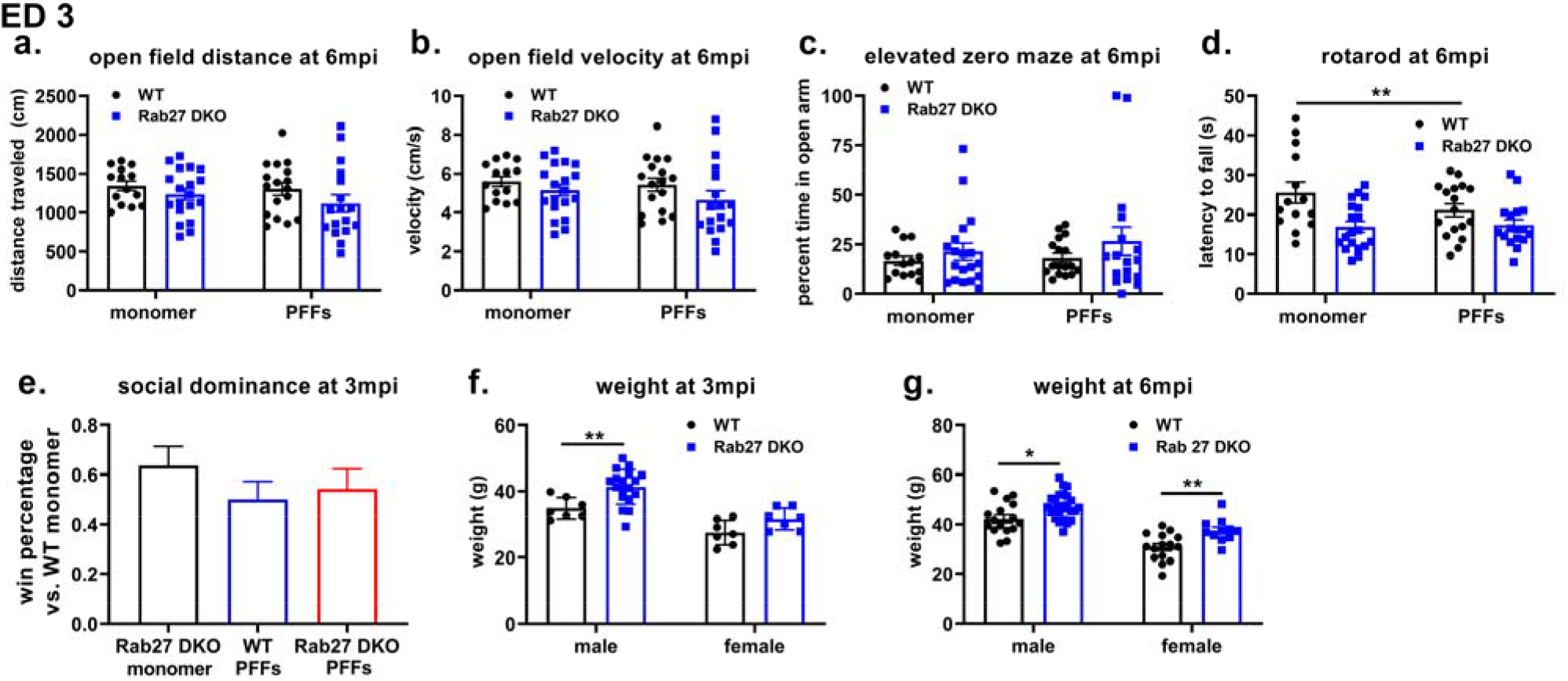
PFF injection does not induce motor or anxiety deficits at 6 months. Total distance traveled (a) and average velocity (b) on the open field test at 6 mpi. n=14-19. c) Percent time spent in the open area in the elevated zero maze at 6 mpi. n=14-19. d) Latency to fall off accelerating rotarod at 6 mpi. n=14-19. e) Win percentage in the social dominance test at 3 mpi. n=9-12. f, g) Mouse weights at 3 mpi (f; n=7-18), and 6 mpi (g; n=11-25). 2-way ANOVA with post-hoc Tukey’s test. *p<0.05, **p<0.01.

## Notes

### Competing Interest Statement

The authors have declared no competing interest.

## REFERENCES

Arimura, N., Kimura, T., Nakamuta, S., Taya, S., Funahashi, Y., Hattori, A., Shimada, A., Menager, C., Kawabata, S., Fujii, K., et al. (2009). Anterograde transport of TrkB in axons is mediated by direct interaction with Slp1 and Rab27. Dev Cell 16, 675–686.

Arrant, A.E., Filiano, A.J., Warmus, B.A., Hall, A.M., and Roberson, E.D. (2016). Progranulin haploinsufficiency causes biphasic social dominance abnormalities in the tube test. Genes, Brain and Behavior 15, 588–603.

Bellomo, G., Paciotti, S., Gatticchi, L., and Parnetti, L. (2020). The vicious cycle between alpha-synuclein aggregation and autophagic-lysosomal dysfunction. Mov Disord 35, 34–44.

Bonet-Ponce, L., and Cookson, M.R. (2019). The role of Rab GTPases in the pathobiology of Parkinson’ disease. Current Opinion in Cell Biology 59, 73–80.

Desnos, C., Schonn, J.-S.b., Huet, S.b., Tran, V.S., El-Amraoui, A., Raposo, G.a., Fanget, I., Chapuis, C., Ménasché, G.l., de Saint Basile, G.v., et al. (2003). Rab27A and its effector MyRIP link secretory granules to F-actin and control their motion towards release sites. Journal of Cell Biology 163, 559–570.

Ding, H., Underwood, R., Lavalley, N., and Yacoubian, T.A. (2015). 14-3-3 inhibition promotes dopaminergic neuron loss and 14-3-3θ overexpression promotes recovery in the MPTP mouse model of Parkinson’s disease. Neuroscience 307, 73–82.

El-Amraoui, A., Schonn, J.S., Küssel-Andermann, P., Blanchard, S., Desnos, C., Henry, J.P., Wolfrum, U., Darchen, F., and Petit, C. (2002). MyRIP, a novel Rab effector, enables myosin VIIa recruitment to retinal melanosomes. EMBO reports 3, 463–470.

Fukuda, M. (2008). Regulation of secretory vesicle traffic by Rab small GTPases. Cell Mol Life Sci 65, 2801–2813.

Fukuda, M. (2013). Rab27 Effectors, Pleiotropic Regulators in Secretory Pathways. Traffic 14, 949–963.

Fukuda, M., Kuroda, T.S., and Mikoshiba, K. (2002). Slac2-a/Melanophilin, the Missing Link between Rab27 and Myosin Va: IMPLICATIONS OF A TRIPARTITE PROTEIN COMPLEX FOR MELANOSOME TRANSPORT. Journal of Biological Chemistry 277, 12432–12436.

Giasson, B.I., Murray, I.V., Trojanowski, J.Q., and Lee, V.M.-Y. (2001). A hydrophobic stretch of 12 amino acid residues in the middle of α-synuclein is essential for filament assembly. Journal of Biological Chemistry 276, 2380–2386.

Ginsberg, S., Mufson, E., Alldred, M., Counts, S., Wuu, J., Nixon, R., and Che, S. (2011). Upregulation of select rab GTPases in cholinergic basal forebrain neurons in mild cognitive impairment and Alzheimer’s disease. Journal of chemical neuroanatomy 42, 102–110.

Gomi, H., Mori, K., Itohara, S., and Izumi, T. (2007). Rab27b Is Expressed in a Wide Range of Exocytic Cells and Involved in the Delivery of Secretory Granules Near the Plasma Membrane. Molecular biology of the cell 18, 4377–4386.

Hayakawa, H., Nakatani, R., Ikenaka, K., Aguirre, C., Choong, C.J., Tsuda, H., Nagano, S., Koike, M., Ikeuchi, T., Hasegawa, M., et al. (2020). Structurally distinct alpha-synuclein fibrils induce robust parkinsonian pathology. Movement disorders : official journal of the Movement Disorder Society 35, 256–267.

Holmes, B.B., DeVos, S.L., Kfoury, N., Li, M., Jacks, R., Yanamandra, K., Ouidja, M.O., Brodsky, F.M., Marasa, J., Bagchi, D.P., et al. (2013). Heparan sulfate proteoglycans mediate internalization and propagation of specific proteopathic seeds. Proceedings of the National Academy of Sciences 110, E3138–E3147.

Kariya, Y., Honma, M., Hanamura, A., Aoki, S., Ninomiya, T., Nakamichi, Y., Udagawa, N., and Suzuki, H. (2011). Rab27a and Rab27b are involved in stimulation-dependent RANKL release from secretory lysosomes in osteoblastic cells. J Bone Miner Res 26, 689–703.

Kimura, T., Kaneko, Y., Yamada, S., Ishihara, H., Senda, T., Iwamatsu, A., and Niki, I. (2008). The GDP-dependent Rab27a effector coronin 3 controls endocytosis of secretory membrane in insulin-secreting cell lines. Journal of Cell Science 121, 3092–3098.

Kimura, T., and Niki, I. (2011). Rab27a, actin and beta-cell endocytosis [Review]. Endocrine Journal 58, 1–6.

Kuroda, T.S., and Fukuda, M. (2005). Functional analysis of Slac2-c/MyRIP as a linker protein between melanosomes and myosin VIIa. Journal of Biological Chemistry 280, 28015–28022.

Leak, R.K., Frosch, M.P., Beach, T.G., and Halliday, G.M. (2019). Alpha-synuclein: prion or prion-like? Acta Neuropathologica 138, 509–514.

Lopes, V.S., Ramalho, J.S., Owen, D.M., Karl, M.O., Strauss, O., Futter, C.E., and Seabra, M.C. (2007). The ternary Rab27a–Myrip–Myosin VIIa complex regulates melanosome motility in the retinal pigment epithelium. Traffic 8, 486–499.

Luk, K.C., Kehm, V., Carroll, J., Zhang, B., O’Brien, P., Trojanowski, J.Q., and Lee, V.M.Y. (2012). Pathological α-synuclein transmission initiates Parkinson-like neurodegeneration in nontransgenic mice. Science (New York, NY) 338, 949–953.

Malik, B.R., Maddison, D.C., Smith, G.A., and Peters, O.M. (2019). Autophagic and endo-lysosomal dysfunction in neurodegenerative disease. Mol Brain 12, 100.

Mao, X., Ou, M.T., Karuppagounder, S.S., Kam, T.-I., Yin, X., Xiong, Y., Ge, P., Umanah, G.E., Brahmachari, S., Shin, J.-H., et al. (2016). Pathological α-synuclein transmission initiated by binding lymphocyte-activation gene 3. Science 353, aah3374.

Miczek, K.A., Brykczynski, T., and Grossman, S.P. (1974). Differential effects of lesions in the amygdala, periamygdaloid cortex, and stria terminalis on aggressive behaviors in rats. J Comp Physiol Psychol 87, 760–771.

Mooren, O.L., Galletta, B.J., and Cooper, J.A. (2012). Roles for Actin Assembly in Endocytosis. Annual Review of Biochemistry 81, 661–686.

Osterberg, Valerie R., Spinelli, Kateri J., Weston, Leah J., Luk, Kelvin C., Woltjer, Randall L., and Unni, Vivek K. (2015). Progressive Aggregation of Alpha-Synuclein and Selective Degeneration of Lewy Inclusion-Bearing Neurons in a Mouse Model of Parkinsonism. Cell Reports 10, 1252–1260.

Ostrowski, M., Carmo, N.B., Krumeich, S., Fanget, I., Raposo, G., Savina, A., Moita, C.F., Schauer, K., Hume, A.N., Freitas, R.P., et al. (2010). Rab27a and Rab27b control different steps of the exosome secretion pathway. Nature Cell Biology 12, 19–30.

Park, J.W., Vahidi, B., Taylor, A.M., Rhee, S.W., and Jeon, N.L. (2006). Microfluidic culture platform for neuroscience research. Nature Protocols 1, 2128–2136.

Pavlos, N.J., Grønborg, M., Riedel, D., Chua, J.J., Boyken, J., Kloepper, T.H., Urlaub, H., Rizzoli, S.O., and Jahn, R. (2010). Quantitative analysis of synaptic vesicle Rabs uncovers distinct yet overlapping roles for Rab3a and Rab27b in Ca2+-triggered exocytosis. Journal of Neuroscience 30, 13441–13453.

Prots, I., Veber, V., Brey, S., Campioni, S., Buder, K., Riek, R., Böhm, K.J., and Winner, B. (2013). α-Synuclein oligomers impair neuronal microtubule-kinesin interplay. J Biol Chem 288, 21742–21754.

Ramalho, J.S., Lopes, V.S., Tarafder, A.K., Seabra, M.C., and Hume, A.N. (2009). Myrip uses distinct domains in the cellular activation of myosin VA and myosin VIIA in melanosome transport. Pigment cell & melanoma research 22, 461–473.

Shen, Y.-T., Gu, Y., Su, W.-F., Zhong, J.-f., Jin, Z.-H., Gu, X.-S., and Chen, G. (2016). Rab27b is Involved in Lysosomal Exocytosis and Proteolipid Protein Trafficking in Oligodendrocytes. Neuroscience Bulletin 32, 331–340.

Shrivastava, A.N., Redeker, V., Fritz, N., Pieri, L., Almeida, L.G., Spolidoro, M., Liebmann, T., Bousset, L., Renner, M., Léna, C., et al. (2015). α-synuclein assemblies sequester neuronal α3-Na+/K+-ATPase and impair Na+ gradient. Embo j 34, 2408–2423.

Soumiya, H., Godai, A., Araiso, H., Mori, S., Furukawa, S., and Fukumitsu, H. (2016). Neonatal Whisker Trimming Impairs Fear/Anxiety-Related Emotional Systems of the Amygdala and Social Behaviors in Adult Mice. PloS one 11, e0158583–e0158583.

Spillantini, M.G., Schmidt, M.L., Lee, V.M.Y., Trojanowski, J.Q., Jakes, R., and Goedert, M. (1997). α-Synuclein in Lewy bodies. Nature 388, 839–840.

Stoyka, L.E., Arrant, A.E., Thrasher, D.R., Russell, D.L., Freire, J., Mahoney, C.L., Narayanan, A., Dib, A.G., Standaert, D.G., and Volpicelli-Daley, L.A. (2020). Behavioral defects associated with amygdala and cortical dysfunction in mice with seeded α-synuclein inclusions. Neurobiology of Disease 134, 104708.

Tolmachova, T., Åbrink, M., Futter, C.E., Authi, K.S., and Seabra, M.C. (2007). Rab27b regulates number and secretion of platelet dense granules. Proceedings of the National Academy of Sciences 104, 5872–5877.

Underwood, R., Wang, B., Carico, C., Whitaker, R.H., Placzek, W.J., and Yacoubian, T.A. (2020). The GTPase Rab27b regulates the release, autophagic clearance, and toxicity of alpha-synuclein. Journal of Biological Chemistry.

Vidyadhara, D.J., Lee, J.E., and Chandra, S.S. (2019). Role of the endolysosomal system in Parkinson’s disease. J Neurochem 150, 487–506.

Volpicelli-Daley, L.A., Luk, K.C., and Lee, V.M. (2014). Addition of exogenous alpha-synuclein preformed fibrils to primary neuronal cultures to seed recruitment of endogenous alpha-synuclein to Lewy body and Lewy neurite-like aggregates. Nat Protoc 9, 2135–2146.

Volpicelli-Daley, L.A., Luk, K.C., Patel, T.P., Tanik, S.A., Riddle, D.M., Stieber, A., Meaney, D.F., Trojanowski, J.Q., and Lee, V.M. (2011). Exogenous alpha-synuclein fibrils induce Lewy body pathology leading to synaptic dysfunction and neuron death. Neuron 72, 57–71.

Wang, B., Underwood, R., Kamath, A., Britain, C., McFerrin, M.B., McLean, P.J., Volpicelli-Daley, L.A., Whitaker, R.H., Placzek, W.J., Becker, K., et al. (2018). 14-3-3 Proteins Reduce Cell-to-Cell Transfer and Propagation of Pathogenic α-Synuclein. The Journal of Neuroscience 38, 8211–8232.

Yamaoka, M., Ando, T., Terabayashi, T., Okamoto, M., Takei, M., Nishioka, T., Kaibuchi, K., Matsunaga, K., Ishizaki, R., Izumi, T., et al. (2016). PI3K regulates endocytosis after insulin secretion by mediating signaling crosstalk between Arf6 and Rab27a. Journal of Cell Science 129, 637.

Yi, Z., Yokota, H., Torii, S., Aoki, T., Hosaka, M., Zhao, S., Takata, K., Takeuchi, T., and Izumi, T. (2002). The Rab27a/granuphilin complex regulates the exocytosis of insulin-containing dense-core granules. Mol Cell Biol 22, 1858–1867.

Zhang, Q., Xu, Y., Lee, J., Jarnik, M., Wu, X., Bonifacino, J.S., Shen, J., and Ye, Y. (2020). A myosin-7B–dependent endocytosis pathway mediates cellular entry of α-synuclein fibrils and polycation-bearing cargos. Proceedings of the National Academy of Sciences 117, 10865–10875.

Zhou, T., Sandi, C., and Hu, H. (2018). Advances in understanding neural mechanisms of social dominance. Current Opinion in Neurobiology 49, 99–107.

## Method References

18 Volpicelli-Daley, L. A. et al. Exogenous alpha-synuclein fibrils induce Lewy body pathology leading to synaptic dysfunction and neuron death. Neuron 72, 57–71, doi:S0896-6273(11)00844-0 [pii] 10.1016/j.neuron.2011.08.033 (2011).

19 Luk, K. C. et al. Pathological α-synuclein transmission initiates Parkinson-like neurodegeneration in nontransgenic mice. Science (New York, N.Y.) 338, 949–953, doi:10.1126/science.1227157 (2012).

44 Tolmachova, T., Åbrink, M., Futter, C. E., Authi, K. S. & Seabra, M. C. Rab27b regulates number and secretion of platelet dense granules. Proceedings of the National Academy of Sciences 104, 5872–5877, doi:10.1073/pnas.0609879104 (2007).

45 Wang, B. et al. 14-3-3 Proteins Reduce Cell-to-Cell Transfer and Propagation of Pathogenic α-Synuclein. The Journal of Neuroscience 38, 8211–8232, doi:10.1523/jneurosci.1134-18.2018 (2018).

46 Giasson, B. I., Murray, I. V., Trojanowski, J. Q. & Lee, V. M.-Y. A hydrophobic stretch of 12 amino acid residues in the middle of α-synuclein is essential for filament assembly. Journal of Biological Chemistry 276, 2380–2386 (2001).

47 Ding, H., Underwood, R., Lavalley, N. & Yacoubian, T. A. 14-3-3 inhibition promotes dopaminergic neuron loss and 14-3-3θ overexpression promotes recovery in the MPTP mouse model of Parkinson’s disease. Neuroscience 307, 73–82 (2015).

